# Gold Nanobipyramids as Second Near Infrared Optical Coherence Tomography Contrast Agents for Multiplexed *In Vivo* Lymphangiography

**DOI:** 10.1101/656744

**Authors:** Peng Si, Saba Shevidi, Edwin Yuan, Ke Yuan, Ziv Lautman, Stefanie S. Jeffrey, George W. Sledge, Adam de la Zerda

## Abstract

Developing contrast-enhanced optical coherence tomography (OCT) techniques is important for specific imaging of tissue lesions, molecular imaging, cell-tracking, and highly sensitive microangiography and lymphangiography. Multiplexed OCT imaging in the second near infrared (NIR-II) window is highly desirable since it allows simultaneous imaging and tracking of multiple biological events in high resolution with deeper tissue penetration *in vivo*. Here we demonstrate that gold nanobipyramids can function as OCT multiplexing contrast agents, allowing the visualization of two separate lymphatic flows occurring simultaneously from different drainage basins into the same lymph node in a live mouse. Contrast-enhanced multiplexed lymphangiography of a melanoma tumor *in vivo* shows that the peritumoral lymphatic drainage upstream of the tumor is unidirectional, with some drainage directly into the tumor, but the lymphatic drainage from the tumor is multi-directional. We also demonstrate real-time tracking of the contrast agents draining from a melanoma tumor specifically to the sentinel lymph node of the tumor and the three-dimensional distribution of the contrast agents in the lymph node.

## Introduction

Optical coherence tomography (OCT) is an emerging imaging modality to investigate tissue microstructural morphology. It allows three-dimensional real time *in vivo* imaging at cellular-scale resolution and several-millimeter tissue penetration depth.^1^ However, since most tissue, cells and biomolecules lack intrinsic OCT contrast, it is necessary to develop contrast-enhanced OCT techniques to investigate the functional processes of the living body, such as growth of lesioned tissue, cell dynamics, and flows in vasculature.^2,3^ Thus far, quite a few materials have been studied as exogenous OCT contrast agents, including microspheres,^4,5^ microbubbles,^6,7^ magnetic nanoparticles^8^ and plasmonic nanoparticles.^9–12^ Among them, plasmonic gold nanoparticles are of particular interest due to their highly tunable shape, size, optical properties, facile surface chemistry and excellent biocompatibility.^13^ Nevertheless, very few reported OCT contrast agents have shown the capability of multiplexing, especially in the second near infrared (NIR-II) window (1100–1400 nm), which allows deeper tissue penetration.^14^ Recently, we reported that gold nanoprisms can be used as OCT contrast agents in the NIR-II window due to their strong scattering at 1385 nm.^12^ In this work, we demonstrate that gold nanobipyramids (GNBPs) with two distinct aspect ratios and surface plasmon resonances can work as OCT contrast agents in the NIR-II window for multiplexed imaging. We show that by subcutaneously injecting two different types of GNBPs, GNBP-I and GNBP-II, two separate lymphatic flows can be visualized simultaneously in a live mouse ear using a custom dual-band OCT spectral signal processing algorithm described in our previous work.^11^ The contrast agents also allow us to map lymphatic flows from two different drainage basins-tumor and peritumoral tissues into the same lymph node. By injecting GNBP-I intratumorally and GNBP-II subcutaneously, the tumor and peritumoral lymphatic drainage pathways can be visualized simultaneously. Our multiplexed imaging shows that a melanoma tumor implanted in the mouse ear drains the lymphatic fluid not only in the cervical direction, but also in all other directions. Subcutaneously injected contrast agents appear not only in the peritumoral lymphatic vessels, but also inside the tumor. We also show that after injection, we can track the contrast agents in different organs of the body. *In vivo* and *ex vivo* lymph node imaging show that the peritumoral lymphatic vessels drain the contrast agents to the deep cervical lymph nodes.

## Results and discussion

GNBPs were prepared according to the method described in Supporting Information. The size and surface morphology of the as-synthesized GNBPs were studied by transmission electron microscopy (TEM). GNBP-I and GNBP-II show similar surface morphology, but different dimensions and aspect ratios (Figure 1a, b). GNBP-I has an average length of 137 nm and width of 22 nm, while GNBP-II is larger with an average length of 177 nm and width of 25 nm. These dimensions correspond to aspect ratios of 6.2 and 7.1 for GNBP-I and GNBP-II, respectively (Figure 1c). The NIR spectra show that the surface plasmon resonances of GNBP-I and GNBP-II are at 1225 nm and 1415 nm, respectively (Figure 1d).

**Figure 1.**
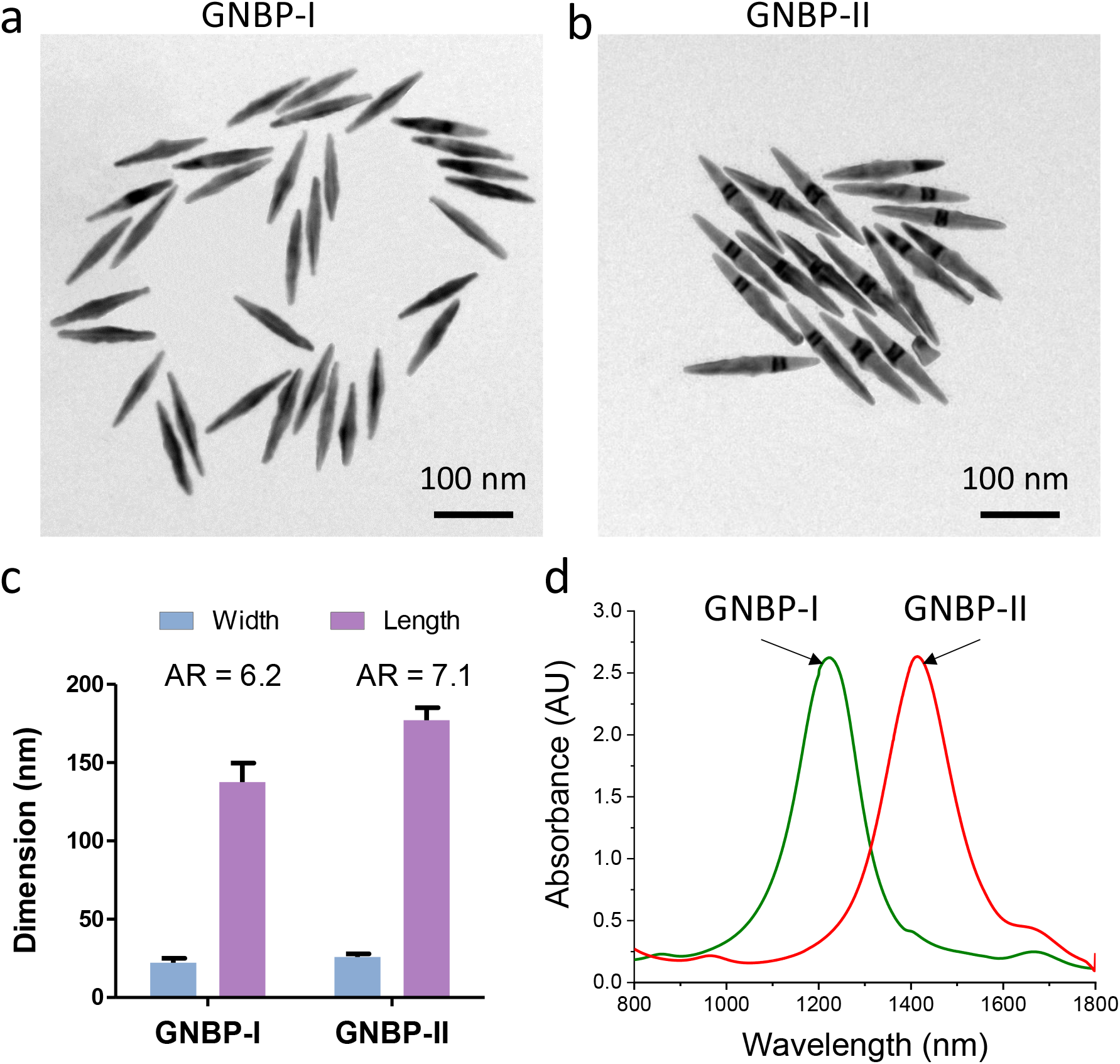
Characterization of GNBPs. (a, b) TEM images of GNBP-I and GNBP-II. (c) The dimensions and aspect ratio (AR) of GNBP-I and GNBP-II. Dimensions are presented as mean ± SEM. (d) The near infrared spectra of GNBP-I and GNBP-II.

To take advantage of the unique spectral characteristics of GNPB-I and GNBP-II, we use a custom dual band spectral analysis algorithm^11^ to detect the nanoparticles. The algorithm divides the OCT interferogram in the source bandwidth into two sub-bands: Band I (1190–1271 nm) and Band II (1272–1365 nm) (see method section in Supporting Information). The interferogram in each sub-band is reconstructed independently to create two OCT images in the spectral domain. The Band II OCT image is then subtracted from the Band I OCT image to obtain the spectral contrast signal. To measure the spectral contrast signal intensity as a function of the concentration of each type of GNBP, we imaged GNBP-I and GNBP-II with a broad range of concentrations in capillary tubes using mouse whole blood as reference. As expected, we observe two distinct spectral contrast signals for GNBP-I and GNBP-II in the reconstructed spectral OCT B-scan images color-coded by a hue-saturation-value (HSV) scheme (Figure 2a, b). GNBP-I shows positive spectral contrast signals since it has enhanced scattering in Band I but reduced scattering in Band II, while GNPB-II shows negative spectral contrast signals because of its opposite response to the two sub-bands. With increasing concentrations, stronger OCT spectral contrast signals can be observed for both types of GNBP. At high concentrations (*e.g.*, 1 nM), the opposite spectral contrast signal can be observed deep in the capillary tube due to a spectral shadowing effect. For quantitative analysis, regions of interest (ROIs) were selected from the top of the tube to avoid the spectral shadowing effect. The analysis shows that the spectral contrast signal of GNBP-I increases proportionally with the nanoparticle concentration from 5 pM to 1 nM (Figure 2c), while the spectral contrast signal of GNBP-II increases linearly in the concentration range of 5 to 100 pM. The spectral contrast signal of GNBP-II can be observed to saturate when the nanoparticle concentration exceeds 500 pM (Figure 2d). We calculate the detection limits of GNBP-I and GNBP-II to be 3.6 and 2.8 pM, respectively.

**Figure 2.**
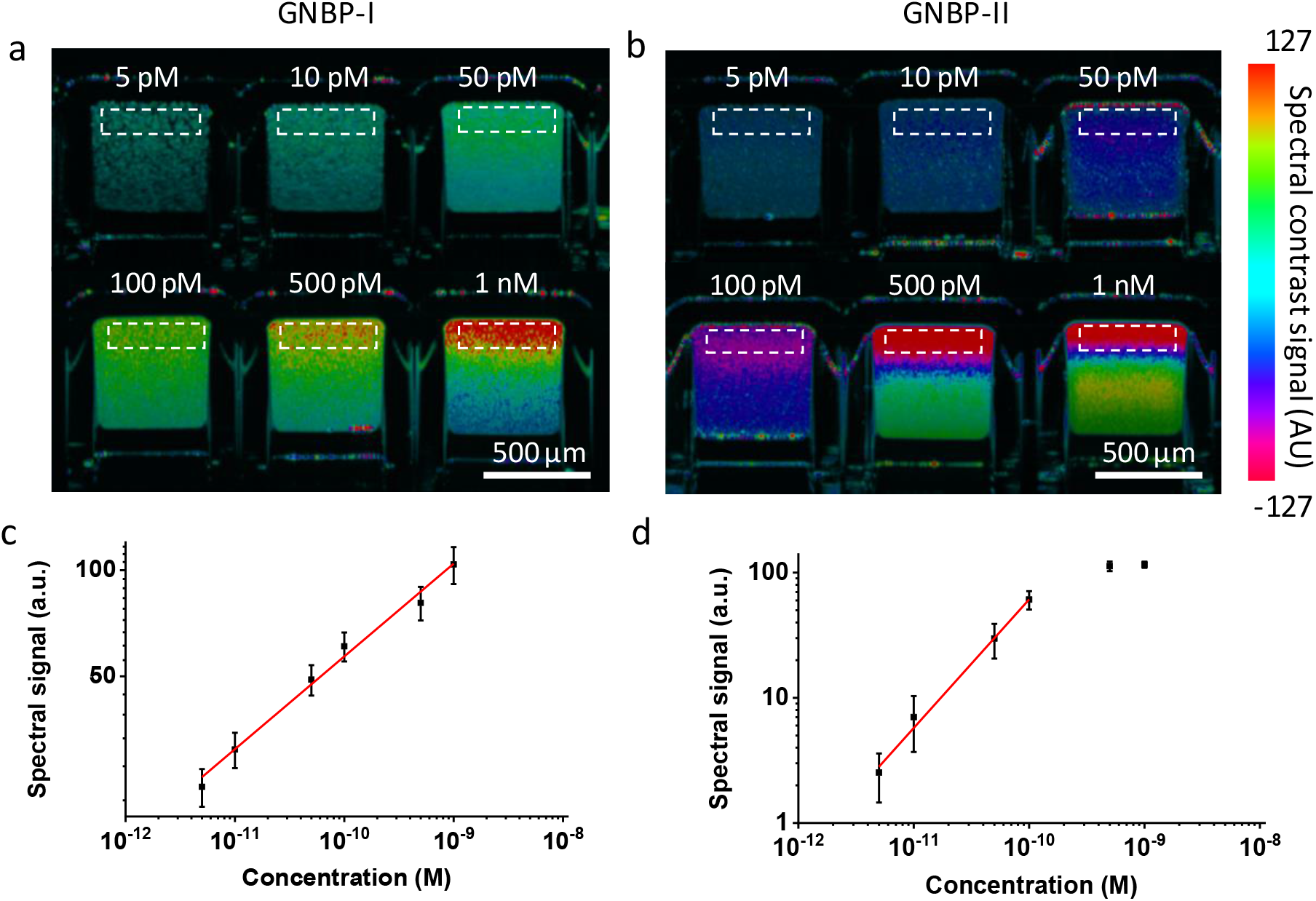
*In vitro* characterization of OCT spectral signals of GNBPs in glass capillary tube phantoms. (a, b) OCT spectral signals of GNBP-I (a) and GNBP-II (b) at different concentrations. The white boxes show the regions of interest (ROIs) for quantitative OCT spectral signal analysis. The ROIs were selected from the top of the tubes because the signal from further down suffers from spectral shadowing. (c, d) Mean OCT spectral signal of GNBP-I (c) and GNBP-II (d) as a function of GNBP concentration in water (n=3 measurements per concentration; data are presented as mean ± SEM).

For *in vivo* experiments, the GNBPs were functionalized with polyethylene glycol (PEG, MW ~5 kDa) to increase their stability and biocompatibility in the biological tissue. To characterize the OCT spectral contrast signals of GNBPs *in vivo*, we subcutaneously injected varying concentrations of PEGylated GNBP-I and GNBP-II in two different mouse ears, and imaged the tissue at each injection site with OCT right after each administration (Figure 3a, b). To display the spectral contrast signals of GNBPs in the tissue, we created cross-sectional compound images by combining OCT structure, spectral contrast, and flow information in an HSV scheme. Stronger OCT spectral contrast signals can be observed in the mouse ear tissues for both GNBPs as the concentrations of the injections are increased (Figure 3c, d). The OCT spectral contrast signals in the mice pinnae increases linearly for GNBP-I concentrations from 5–50 pM (Figure 3e) and for GNBP-II from 5–100 pM (Figure 3f).

**Figure 3.**
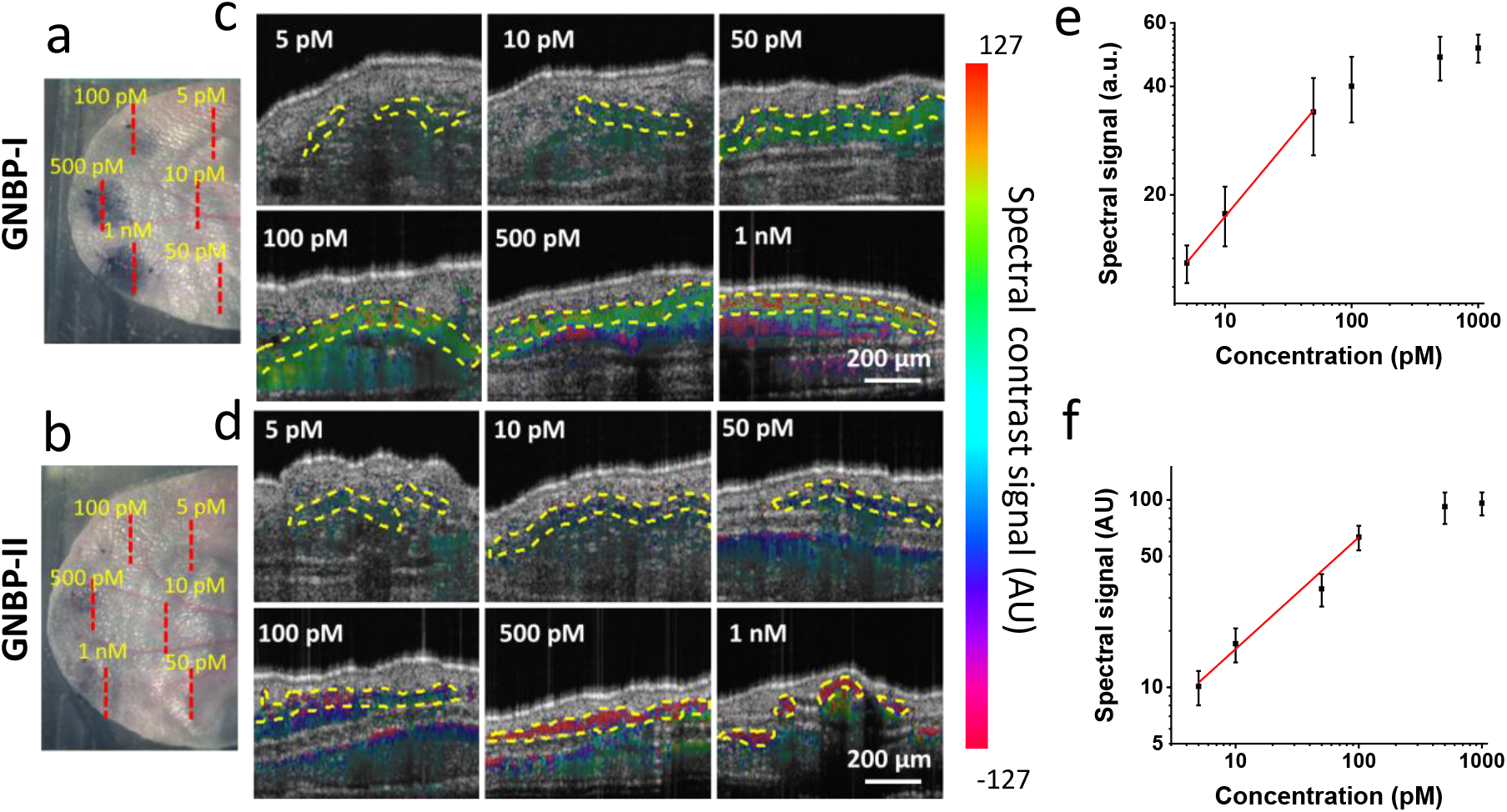
*In vivo* characterization of OCT spectral contrast signals of GNBPs in mouse pinnae. (a, b) Photographs of mouse ears after being injected subcutaneously with different concentrations of GNBP-I (a) and GNBP-II (b). Red dashed lines indicate locations of the cross-sectional scans presented in (c, d). (c, d) Cross-sectional compound OCT images of mouse ears with different concentrations of GNBP-I (c) and GNBP-II (d) injected in the tissue. The compound images are created by combining OCT structure and flow-gated spectral contrast signals in an HSV scheme. The areas enclosed by yellow dashed lines are used as ROIs to quantify the OCT spectral signals. The ROIs are selected as the top 30 pixels that pass the flow gate for each A-scan or all such pixels when there are fewer than 30 of them for an A-scan. (e, f) Mean OCT spectral signals of GNBP-I (e) and GNBP-II (f) in the tissue as a function of the injected GNBP concentration (n=3 mouse measurements per concentration; data are presented as mean ± SEM).

To image the two contrast agents simultaneously *in vivo*, we subcutaneously injected PEGylated GNBP-I and GNBP-II in two separate distal locations on a mouse ear in a sequential manner (Figure 4a-c). We imaged the mouse ear before and after each injection, and produced “flow-gated” spectral OCT images using our dual band signal processing algorithm (See Methods) to show the spectral contrast signals only in the blood and lymphatic vessels. (Figure 4d-f). At pre-injection, the flow-gated OCT spectral image is an angiogram which shows only the network of blood vessels (Figure 4d), since the lymphatic fluid is optically transparent and thus has no signal in the flow-gated image. The spectral contrast signals in this image are close to zero (color coded as cyan) because blood has almost equal signals in each of the two sub-bands of our spectral reconstruction algorithm. In a cross-sectional compound image (Figure 4g), one major blood vessel (indicated by a red arrow) and three lymphatic vessels (yellow arrows) can be visualized. The lymphatic vessels appear dark because lymphatic fluid produces negligible optical reflection.^15^ Upon subcutaneously administering GNBP-I, we observed an extensive network of vasculature with high spectral contrast signals in addition to the blood vessels on the flow-gated OCT spectral image (Figure 4e). These vasculatures are presumably lymphatic vessels which drain the injected contrast agents from the interstitial tissue. The cross-sectional compound image (Figure 4h) shows that all three lymphatic vessels exhibit high spectral contrast signals (arrows), confirming that the spectrally positive network that appeared on the post-injection flow-gated OCT spectral image are indeed lymphatic vessels. 30 min after the first injection, we subcutaneously injected the second contrast agent, PEGylated GNBP-II, at a different distal location of the ear (Figure 4c). After this second injection, negative spectral contrast signals can be observed in some lymphatic vessels on the flow-gated OCT spectral image (red arrows in Figure 4f). The spectral contrast signals of some other lymphatic vessels remain positive but show reduced spectral intensity (*e.g.*, white arrow). The reduced spectral intensity could be attributed either to decreased concentration of GNBP-I due to lymphatic clearance over time, or to spectral contrast signal neutralization by an influx of GNBP-II. In the cross-sectional compound image (Figure 4i), negative spectral contrast signals can be observed in one of the lymphangia (yellow arrow). On close inspection of the lymphangiogram, we can visualize some junctions that separate two distinct spectral contrast signals in the lymphatic vessels (Figure S1). These junctions may correspond to the lymphatic valve structures present at each end of the lymphangion compartments, which maintain the unidirectional flow of lymphatic fluid.^16^

**Figure 4.**
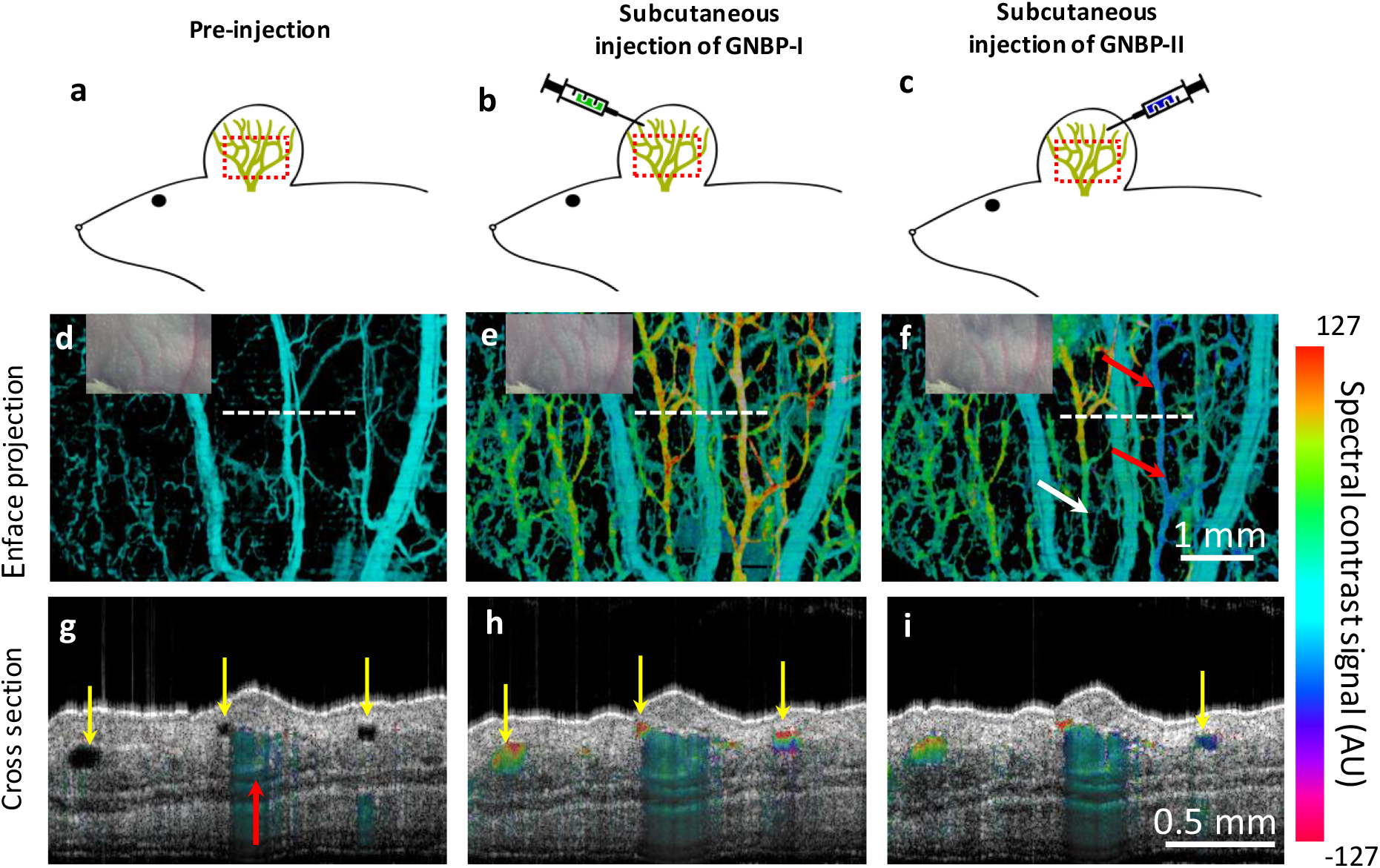
*In vivo* multiplexed lymphangiography of GNBP-I and GNBP-II. (a-c) Schematic illustration of pre-injection stage (a), subcutaneous injection of GNBP-I (b), and subcutaneous injection of GNBP-II (c). The red boxes represent the field of view of the en face OCT images. The green networks represent lymphatic vessels. (d-f) En face flow-gated spectral OCT image of the mouse ear before injection (d), after subcutaneous injection of GNBP-I (e), and after subsequent subcutaneous injection of GNBP-II (f). The insets show photographs of the mouse ear at each imaging stage. The dashed lines show the locations of the cross-sectional images in (g-i). Red arrows in (f) indicate vessels with negative spectral contrast signal characteristic of GNBP-II. White arrow indicates a lymphatic vessel with a reduced positive spectral contrast signal compared to (e). (g-i) Cross-sectional compound OCT images of the mouse ear before injection (g), after GNBP-I injection (h), and after subsequent GNBP-II injection (i). Red arrow indicates a major blood vessel. Yellow arrows indicate lymphatic vessels, all of which have high positive spectral contrast signals in (h), and one of which has a negative spectral contrast signal in (i).

To image tumor lymphatic drainage, we injected PEGylated GNBP-I intratumorally followed by subcutaneous injection of PEGylated GNBP-II upstream of the tumor (Figure 5a-c). OCT B-scan images taken during the intratumoral injection confirm the needle tip is positioned in the middle of the tumor (Figure S2). The angiogenic tumoral and peritumoral blood vessels can be visualized in the pre-injection OCT angiogram (Figure 5d). These vasculatures can also be visualized in the cross-sectional compound image (Figure 5g). After the intratumoral injection, positive spectral contrast signals can be observed in the entire tumor due to the diffusion of GNBP-I throughout the tumor (Figure 5e, h). The lymphatic drainage pathways of the intratumorally injected contrast agents can be readily visualized from the enface flow-gated OCT spectral image. Positive spectral contrast signals can be observed not only in the cervical-direction lymphatic vessels connecting to the tumor, but also in peritumoral lymphatic vessels traveling in various other directions (white arrows in Figure 5e). In a separate study, we show further evidence that the tumor drains the lymphatic fluid in all directions, in contrast to the unidirectional lymphatic flow (cervical direction only) in normal tissue (Figure S3). The multi-directional lymphatic drainage could be caused by malfunctional lymphatic valves in the peritumoral lymphatic vessels.^17^ We also show that the peritumoral lymphatic vessels have larger average vascular diameters compared to normal lymphatic vessels (Figure S4), possibly resulting from hyperplasia.^17,18^

**Figure 5.**
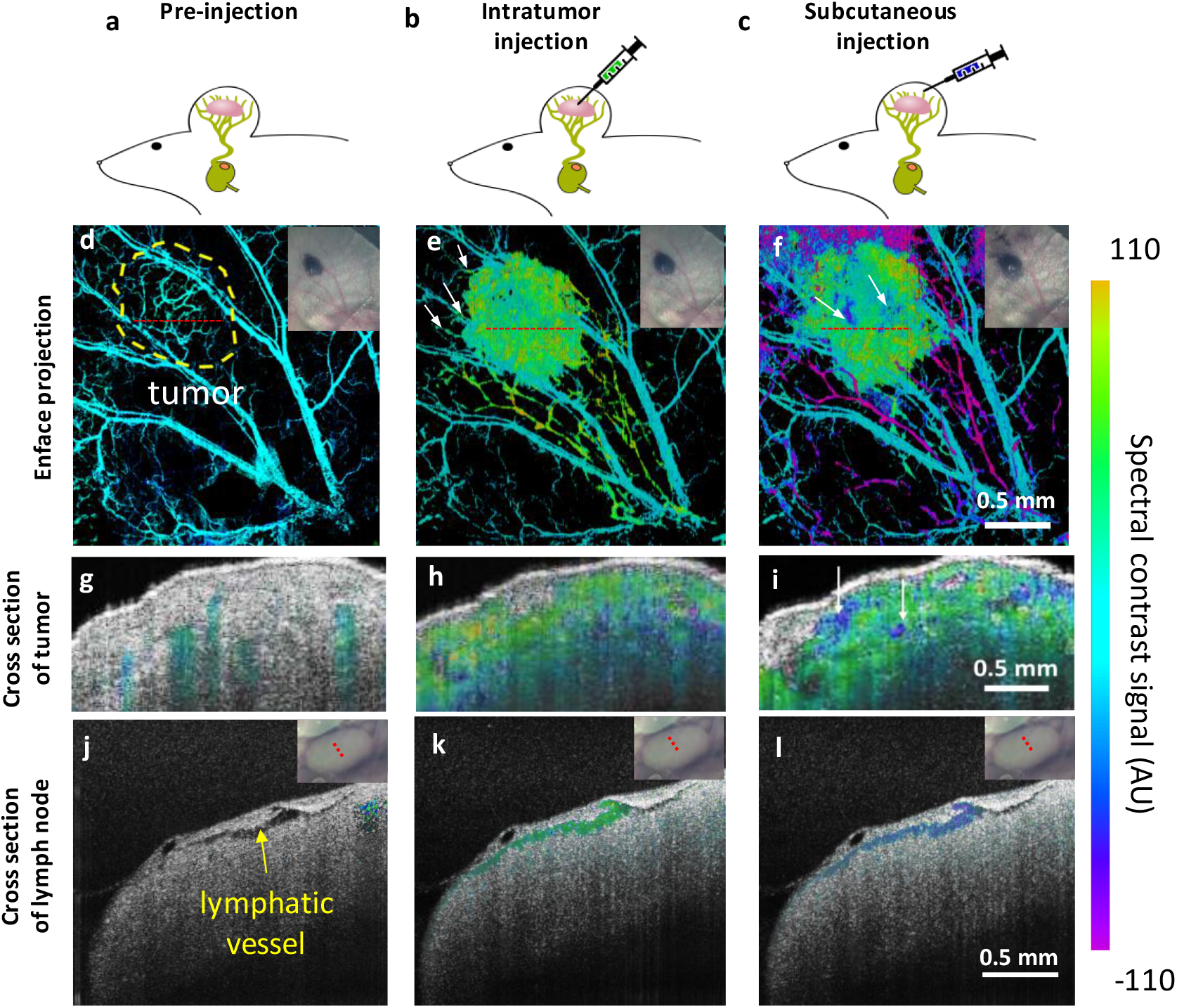
*In vivo* multiplexed imaging of tumor and peritumoral lymphatic drainage. (a-c) Schematic illustration of pre-injection stage (a), intratumor injection of GNBP-I (b), and subsequent subcutaneous injection of GNBP-II (c). The lymphatic vessels are shown draining to a deep cervical lymph node (green). (d-f) En face flow-gated spectral OCT images of the mouse ear before injection, after intratumor injection of GNBP-I, and after subsequent subcutaneous injection of GNBP-II. The dashed circle in (d) marks the primary tumor. The arrows in (e) indicate lymphatic flow in the non-cervical direction. The arrows in (f) point to spectral contrast signals characteristic of GNBP-II, which show that it has flowed into the tumor after the subcutaneous injection. The red dashed lines in (d-f) show the locations of cross-sectional images in (g-i). Insets are photos of the ear near the tumor at each stage. (g-i) Cross-sectional compound images of the mouse ear before injection, after intratumor injection, and after subcutaneous injection. The arrows in (i) indicate spectral contrast signals characteristic of GNBP-II in the tumor. (j-l) *In vivo* OCT cross-sectional compound images of the ipsilateral deep cervical lymph node before injection, 30 min after intratumor injection, and 30 min after subcutaneous injection of the ear. Insets show photos of the lymph node with the cross-sectional location marked by red dashed line. The spectral contrast signal of each contrast agent is clearly visible in the respective compound images (k) and (l).

Following subcutaneous injection of PEGylated GNBP-II, we noticed that negative spectral contrast signals can be visualized not only in peritumoral lymphatic vessels (Figure 5f), but also in the tumor (arrows in Figure 5f, i). This result implies that the subcutaneously injected contrast agents infiltrated into the tumor lymphatic vessels. The cross-sectional compound image reveals that GNBP-II appears in both superficial and deep tumor lymphangia, which are measured to be 50 and 160 μm below the skin surface (arrows in Figure 5i). The superficial lymphangion could correspond to lymphatic vessels in the dermis tissue surrounding the melanoma tumor. However, the lymphangion located at 160 μm below the skin surface corresponds to a tumor lymphatic vessel. The peritumoral lymphatic vessels downstream of the tumor mainly show negative spectral contrast signals after subcutaneous injection of GNBP-II, possibly as a result of reduced concentration of GNBP-I in those vessels due to lymphatic clearance. The tumor and peritumoral lymphatic drainage pathways on the mouse ear are further illustrated schematically in Figure S5.

To further study the lymphatic drainage pathways of the contrast agents injected into the two different basins, we imaged the deep cervical lymph node on the ipsilateral side of the mouse ear pre-injection and after each injection. At pre-injection, no OCT signals can be observed in the lymphatic vessels of the lymph node on the cross-sectional compound image (arrow in Figure 5j). 30 min after intratumoral injection, positive spectral contrast signals can be observed in the lymphatic vessel in the lymph node (Figure 5k). Thirty min after subcutaneous injection, negative spectral contrast signals can be observed in the same lymphatic vessel (Figure 5l). The result corroborates that after intratumoral and subcutaneous injection, each contrast agent was drained into the peritumoral lymphatic vessels and then collected by the deep cervical lymph node.

After the *in vivo* imaging of the melanoma tumor-implanted mouse, we resected deep cervical lymph nodes on both the ipsilateral and contralateral sides (Figure 6, a-c) and conducted three-dimensional (3-D) lymphangiography *ex vivo*. In the spectral 3-D OCT lymphangiograms and cross-sectional compound images, the spectral contrast signals can be visualized in the lymphatic vessels of the ipsilateral cervical lymph node (Figure 6d,e, Movie S1), but not in the contralateral lymph node of the mouse (Figure 6f,g, Movie S2). The lymphatic vessel network and lymphatic follicles in the ipsilateral cervical lymph node can be clearly visualized from the 3-D spectral contrast-enhanced OCT lymphangiogram (Figure S6d and Movie S3). In other words, we have tracked the lymphatic drainage of the contrast agents specifically to the sentinel lymph node of the melanoma tumor.

**Figure 6.**
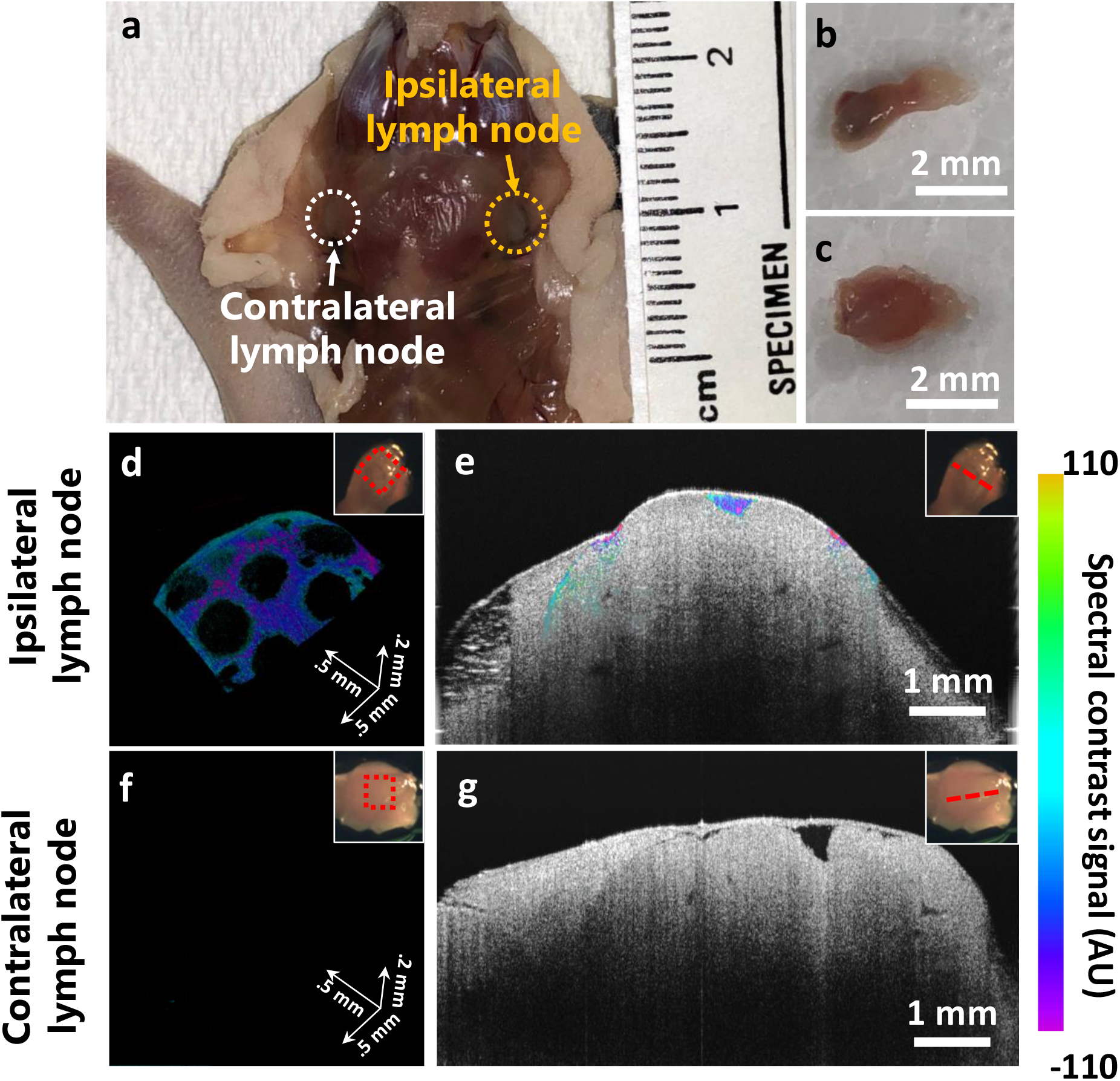
*Ex vivo* imaging of contrast agents in deep cervical mouse lymph nodes. (a-c) Photographs of the deep cervical lymph nodes on the melanoma-implanted mouse (a) and the resected ipsilateral (b) and contralateral (c) deep cervical lymph nodes. (d, e) 3-D OCT lymphangiogram (d) and cross-sectional compound OCT image (e) of the ipsilateral lymph node. Spectral contrast signals of GNBP-II are clearly visible. (f, g) 3-D OCT lymphangiogram (f) and cross-sectional compound OCT image (g) of the contralateral lymph node. No spectral contrast signals can be visualized. Insets are photos of the lymph nodes with the positions of the OCT images marked in red. Movies S1-S3 available in Supplementary Information show fly-through videos of cross-sectional compound images containing (e) and (g) and a rotating 3-D OCT lymphangiogram of (d).

## Conclusion

We have developed a class of spectrally identifiable OCT contrast agents-GNBPs and demonstrated that PEGylated GNBPs can be used for OCT multiplexing in live animals in the NIR-II window. The two types of spectrally distinct contrast agents: GNBP-I and GNBP-II, can be visualized simultaneously in live mouse tissue. By injecting PEGylated GNBP-I and GNBP-II separately in the tumor and in subcutaneous tissue upstream of the tumor, the lymphatic drainage pathways of the tumor and peritumoral tissue were revealed. The multiplexed imaging showed that the tumor drained the contrast agents multi-directionally, while peritumoral tissue drained the contrast agents into the tumor and peritumoral lymphatic vessels. The tumor and peritumoral lymphatic drainage pathways were further illuminated by *in vivo* lymph node imaging after each injection. The OCT spectral contrast signals of both contrast agents can be visualized in the same deep cervical lymph node 30 min after injection. Furthermore, we obtained a spectral 3-D lymphangiogram of the lymph node by imaging the resected lymph node after sacrificing the mouse. The network of lymphatic vessels and lymphatic follicles can be clearly visualized on the 3-D lymphangiogram.

The novel NIR-II OCT contrast agents and multiplexing technique we report here provide a platform that opens many opportunities for biological and clinical research in the future, including multiplexed cell tracking and molecular imaging in live subjects. In the lymphatic research field, this technique could be used for imaging tumor lymph node metastasis of heterogeneous tumor cells (*i.e.*, tumor cells expressing different biomarkers). It could also be employed for molecular imaging to study the dynamic expression of various lymph endothelial cell receptors (*i.e.*, lymphatic vessel endothelial hyaluronan receptor 1 or LYVE-1, and vascular endothelial growth factor receptor 3 or VEGFR-3) in tumor angiogenic lymphatic vessels and lymph nodes. By applying this technique to *in vivo* brain imaging, we believe it can further unravel the mystery and complexity of brain lymphatic drainage pathways, which play important roles in neuron inflammation and neurodegenerative diseases.

## Associated Content

### Supporting Information includes

Description of: preparation and characterization of contrast agents, the OCT instrumentation, *in vitro* imaging, animal handling and *in vivo* imaging, image processing and analysis (Methods); magnified flow-gated OCT spectral image showing lymphatic vessel junctions (Figure S1); photographs and OCT B-scans of intratumoral injection (Figure S2); schematics and flow-gated OCT spectral images comparing lymphatic flow patterns in normal versus tumor lymphatic vessels (Figure S3); flow-gated spectral OCT images and analysis of lymphatic vessel diameters for normal versus peritumoral tissue (Figure S4); schematic illustrations of tumoral and peritumoral lymphatic drainage (Figure S5).

Movie S1: Fly-through video displaying the 3-D compound images of resected deep cervical lymph node on ipsilateral side (30 min after subcutaneous injection) created by combining OCT structure and flow-gated spectral contrast signals in an HSV scheme. (AVI)

Movie S2: Fly-through video displaying the 3-D compound images of resected deep cervical lymph node on contralateral side (30 min after subcutaneous injection) created by combining OCT structure and flow-gated spectral contrast signals in an HSV scheme. (AVI)

Movie S3: Rotating 3-D spectral OCT lymphangiogram showing lymphatic vessel network and lymphatic follicles in the ipsilateral cervical lymph node. (AVI)

## Acknowledgments

This work was funded in part by grants from the United States Air Force (FA9550-15-1-0007), the National Institutes of Health (NIH DP50D012179), the National Science Foundation (NSF 1438340), the Damon Runyon Cancer Research Foundation (DFS# 06-13), Claire Giannini Fund, the Susan G. Komen Breast Cancer Foundation (SAB15-00003), the Mary Kay Foundation (017-14), the Skippy Frank Foundation, the Donald E. and Delia B. Baxter Foundation, a seed grant from the Center for Cancer Nanotechnology Excellence and Translation (CCNE-T; NIH-NCI U54CA151459), and a Stanford Bio-X Interdisciplinary Initiative Seed Grant (IIP6-43). A.d.l.Z is a Chan Zuckerberg Biohub investigator and a Pew-Stewart Scholar for Cancer Research supported by The Pew Charitable Trusts and The Alexander and Margaret Stewart Trust.

